# Adaptations of microbial genomes to human body chemistry

**DOI:** 10.1101/2023.02.12.528246

**Authors:** Jeffrey M. Dick

## Abstract

Water and oxygen availability vary in normal physiology and disease, so evolutionary adjustments of protein sequences to optimally use these chemical resources would represent a competitive advantage for host-associated microbial genomes. In this study, reference proteomes for taxa derived from the Genome Taxonomy Database (GTDB) were combined with 16S rRNA-based taxonomic abundances in order to calculate chemical metrics for community reference proteomes. This permits new insight into community-level genomic adaptation to specific chemical conditions in body sites. Surprisingly, reference proteomes for gut communities appear to be shaped by the physiological function of water absorption in the intestine more than by reducing conditions. Reference proteomes of gut communities in COVID-19 and inflammatory bowel disease (IBD) patients are generally more reduced than controls despite higher relative abundances of aerotolerant organisms and lower abundances of *Faecalibacterium* and other obligate anaerobes. The trend of chemical reduction in patients is supported by multi-omics (i.e., metagenomic and metaproteomic) data for COVID-19 and can be attributed to relatively oxidized protein sequences for obligate anaerobes compared to aerotolerant genera in gut communities. Genomic adaptation to transiently oxygenated conditions, reflected in more oxidized protein sequences, may be an evolutionary strategy for obligate anaerobes to compete with aerotolerant organisms in the chemical context of gut inflammation.

**Impact statement:** How host-associated microbes (the microbiota) interact with body chemistry is important for understanding the chemical factors that may contribute to diseases. Although COVID-19 and IBD are associated with oxidative conditions in the gut due to inflammation, protein sequences inferred for microbial communities in patients exhibit a trend of chemical reduction rather than oxidation. This implies an evolutionary strategy for obligate anaerobes to more effectively compete with oxygen-tolerant organisms that might otherwise dominate the gut during inflammation. The genomes of organisms in gut communities are distinguished from those in other body sites by proteins with lower hydration state, suggesting that physiological gradients of water availability are a major driver of the evolution of human microbiota.

## Introduction

The chemical environment is important for not only free-living microbial communities but also those in host-associated habitats (Pfister et al, 2022). For instance, oxygen concentration differs among body sites in normal conditions and is affected by several diseases (Espey, 2013; Rigottier-Gois, 2013). The gut maintains a steep radial oxygen gradient that may be disrupted in inflammation, promoting the expansion of aerotolerant taxa including facultative anaerobes and microaerophiles (Shin et al, 2015; Lee et al, 2022); such a deviation from normal community composition is known as dysbiosis (Donaldson et al, 2016). Dysbiosis of gut communities is associated with various diseases, including inflammatory bowel disease (IBD) and COVID-19 (Shin et al, 2015; Zhang et al, 2023). However, there is limited information about the genomic adaptation of microbiota to oxidation-reduction (redox) conditions in the gut and other body sites.

Water is a substrate for even more metabolic reactions than oxygen (Frenkel-Pinter et al, 2021). The many roles of water in human physiology are reflected in decreasing organismal water content during development from embryo to adult (Moulton, 1923; Toro-Ramos et al, 2015), higher water content in tumors compared to normal tissue (Dick, 2021; Penet et al, 2021), and gains and loses of H_2_O from eukaryotic cells during transitions to cellular proliferation and dormancy (Munder et al, 2016; Marakhova et al, 2019). Furthermore, a major function of the intestine is absorption of water from chyme (Leiper, 2015), and lower water content in the colon is associated with greater mucosal thickness toward the rectum (Tropini et al, 2017). Therefore, monitoring both oxidation and hydration state may be important for understanding adaptation of microbial genomes to human host habitats.

Host-associated microbiota assemble and mature at timescales of human lives, but microbial lineages have co-evolved with the ancestors of modern humans for millions of years (Ley et al, 2008; Moeller et al, 2016). It follows that the chemical differences of proteins encoded by genomes provide insight into adaptation on evolutionary timescales. Here, the influence of chemical factors on microbial genomic adaptation is explored with a geochemical biology approach. The elemental composition of proteins inferred from multi-omics data is used to calculate two chemical metrics, stoichiometric hydration state (*n*_H2O_) and oxidation state (*n*_O2_ ), which refer to the numbers of water and oxygen molecules in a theoretical reaction to form a protein from thermodynamic components, normalized by the number of amino acid residues (see Methods for details). It was previously proposed that environmental oxygen levels shape the elemental composition of protein sequences during evolution (Acquisti et al, 2007; Vecchio-Pagan et al, 2017); *n*_O2_ is a new metric to test the hypothesis. The utility of protein *n*_H2 O_ as a metric related to environmental variation is supported by previous analyses showing that lower water availability is a selective factor in the evolution and expression of less hydrated proteins as measured by stoichiometric hydration state.

Specifically, *n*_H2O_ decreases with salinity in environmental metagenomes and with hyperglycemic conditions in cell culture and increases in multiple cancer types compared to normal tissue, which suggests a proteomic response to the higher water content of tumors (Dick et al, 2020; Dick, 2021).

Publicly available 16S rRNA datasets were repurposed for a chemical analysis by combining taxonomic abundances with reference proteomes for taxa in order to generate reference proteomes for communities (Dick and Kang, 2023). The community-level trends of chemical metrics produced using this method are consistent with those inferred from metagenomic data for the same samples. Then, community reference proteomes were used to observe patterns of chemical metrics for communities in different body sites and differences between communities associated with inflammatory diseases (COVID-19 and IBD). The main results are that low *n*_H2O_ is the major feature that distinguishes gut communities from other body sites, and that gut communities have more reduced proteins in patients compared to healthy controls. The latter is a surprising result because gut inflammation is commonly associated with oxidative rather than reducing conditions, but this result can be explained by a lower abundance of obligate anaerobes with relatively relatively high-*n*_O2_ reference proteomes during inflammation. Genomic adaptations that code for relatively oxidized proteins may be a new evolutionary mechanism for obligate anaerobes to compete with aerotolerant organisms in the inflammatory gut environment.

## Results

### Consistency between shotgun metagenomes and community reference proteomes

The reliability of chemical differences inferred from community reference proteomes needs to be established by comparison with independent data. For this purpose, we can analyze 16S rRNA sequences and shotgun metagenomes for the same samples available from the Human Microbiome Project (HMP) (The Human Microbiome Project Consortium, 2012). The selected HMP samples include 49 samples analyzed by Aßhauer et al (2015) together with 52 other samples analyzed by Dick and Tan (2023).

The pipeline for processing shotgun metagenomes (hereafter just “metagenomes”) includes a step to screen and remove human DNA by using the bowtie tool and the GRCh38 reference human genome (see Methods for details). In order to assess the contribution of putative human DNA, the pipeline was run twice for each HMP metagenome: once with no screening, and once with screening for human DNA removal. Screening human DNA results in a greater number of samples with low protein prediction rate (i.e., the number of predicted protein sequences as a percentage of the number of analyzed metagenomic reads) (Supplementary file 1). These samples exhibit relatively high scatter of chemical metrics (Figure 1A–B) and may represent low-microbial biomass samples. Because of this, all metagenomic sequencing runs analyzed in this study were subject to a protein prediction rate cutoff of 40%; samples that do not meet this criterion are shown in the plots by open symbols and were omitted from the statistical analysis. Figure 1 shows that screening human DNA from the HMP metagenomes and omitting samples with low protein prediction rate leads to a high correlation of chemical metrics between metagenomically inferred proteins and 16S rRNA-based community reference proteomes. Furthermore, scatter plots of *n*_H2 O_ vs *n*_O2_ show overall consistency between community reference proteomes and proteins inferred from human-DNA-screened metagenomes (Figure 1C).

**Fig. 1.**
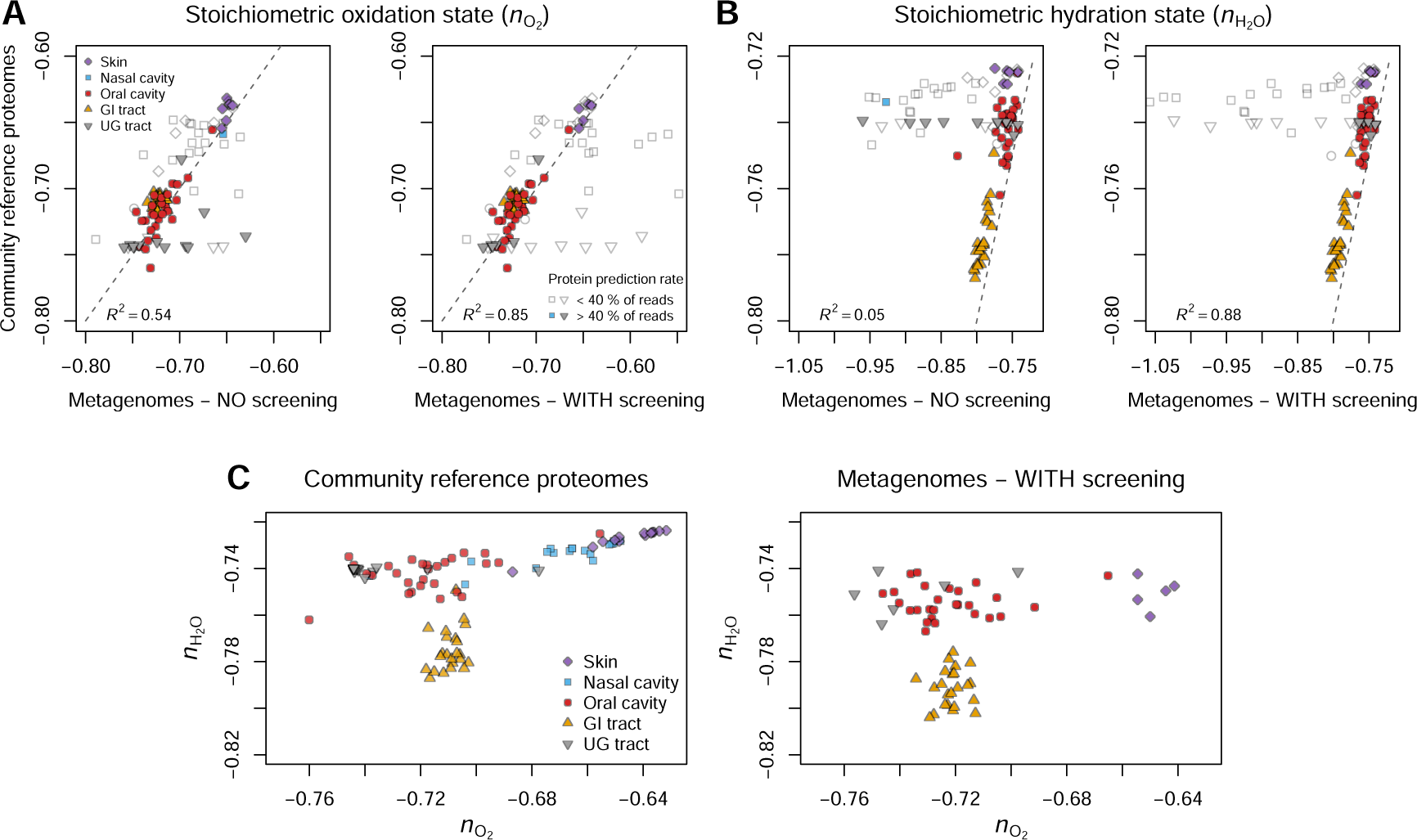
Chemical metrics for community reference proteomes and for proteins inferred from metagenomes for the same samples from the Human Microbiome Project. (**A**) Stoichiometric oxidation state (*n*_O2_ ) and (**B**) stoichiometric hydration state (*n*_H2O_). Chemical metrics were calculated for metagenomes without a screening step to remove human DNA sequences (left plots of A and B) and with the screening step (right plots of A and B). Filled symbols indicate runs for which the number of predicted protein sequences is > 40% of the number of metagenomic reads input to the sequence processing pipeline (see Supplementary file 1); only these samples were used in the calculation of *R*^2^ values. Dashed lines are 1:1 lines, not regression lines. (**C**) Scatter plots of *n*_H2 O_ vs *n*_O2_ for community reference proteomes and metagenomes. Metagenomic processing in the latter plot included the screening step to remove human DNA, and only metagenomic sequencing runs with at least 40% protein prediction rate are shown. GI – gastrointestinal; UG – urogenital. **Figure supplement 1.** Percentages of sequences removed by human DNA screening of HMP metagenomes.

In shotgun metagenomic datasets where there are high levels of human DNA, screening of human DNA may leave few predicted bacterial protein sequences, making estimates of bacterial amino acid composition more uncertain. Specifically, metagenomic samples for nasal cavity and urogenital tract have relatively high amounts of putative human DNA removed in the screening step (Figure 1–figure supplement 1), and none of the screened metagenomic samples for nasal cavity passes the protein-prediction-rate cutoff of 40% (Figure 1C). However, community reference proteomes exhibit relatively high *n*_O2_ for the microbiota of nasal and skin sites. This shows how 16S amplicon sequencing can be used to gain insight into community-level genomic variation in samples where shotgun metagenomes are challenged by high levels of host-derived DNA.

### Assessing possible effects of contamination

The reference proteomes and 16S rRNA training set for taxonomic classification used in this study are based on genomes in the Genome Taxonomy Database (GTDB) that were used without checking for contamination. To increase confidence in chemical metrics for reference proteomes, those generated from GTDB are compared here with contamination-checked genomes available in the MGnify database. Reference proteomes were derived in this study from 65703 genomes in GTDB, representing a total of 16686 genera. Conversely, the Unified Human Gastrointestinal Genome (UHGG v2.0.1) from MGnify (Almeida et al, 2021; Richardson et al, 2023) consists of 4744 high-quality (CheckM contamination < 5% and completeness > 50%) species-level clusters representing 1031 genera. In this version of UHGG, genomes likely to contain chimeric sequences and contigs likely to originate from the host genome were removed (https://ftp.ebi.ac.uk/pub/databases/metagenomics/mgnify_genomes/human-gut/v2.0.1/README_v2.0.1.txt, accessed on 2024-01-02). Taxonomic classifications performed in this study identified 27 genera with high relative abundance changes in gut datasets for COVID-19 and IBD (see Figure 5 below). All but one of these genera is present in the UHGG; the exception is *Pseudocitrobacter*, which is a facultatively anaerobic bacterium isolated from feces (Kämpfer et al, 2014). Therefore, classification of 16S rRNA sequences using the GTDB-based training set successfully identifies gut-associated organisms rather than spurious taxonomic groups, despite the much higher taxonomy diversity of GTDB compared to UHGG.

Species-level reference proteomes in GTDB form natural clusters on a *n*_H2 O_–*n*_O2_ plot (Figure 2A), suggesting that chemical metrics for genus reference proteomes represent genuine biological differences. Similar clusters characterize reference proteomes of taxa selected from UHGG using stringent criteria (contamination < 2% and completeness > 95%, including 2350 species-level clusters representing 643 genera), and chemical metrics for genus-level reference proteomes are highly correlated between GTDB and UHGG (Figure 2B–C). Furthermore, for HMP samples, there is a tight correlation of chemical metrics calculated from the GTDB-based training set and reference proteomes used in this study and from the Ribosomal Database Project (RDP) training set and RefSeq-based reference proteomes used in a previous study (Dick and Tan, 2023) (Figure 2D). Therefore, in comparison to the RDP training set, any classification errors which may specific to the GTDB-based 16S rRNA training set do not adversely affect the calculation of community-level chemical metrics for these samples.

**Fig. 2.**
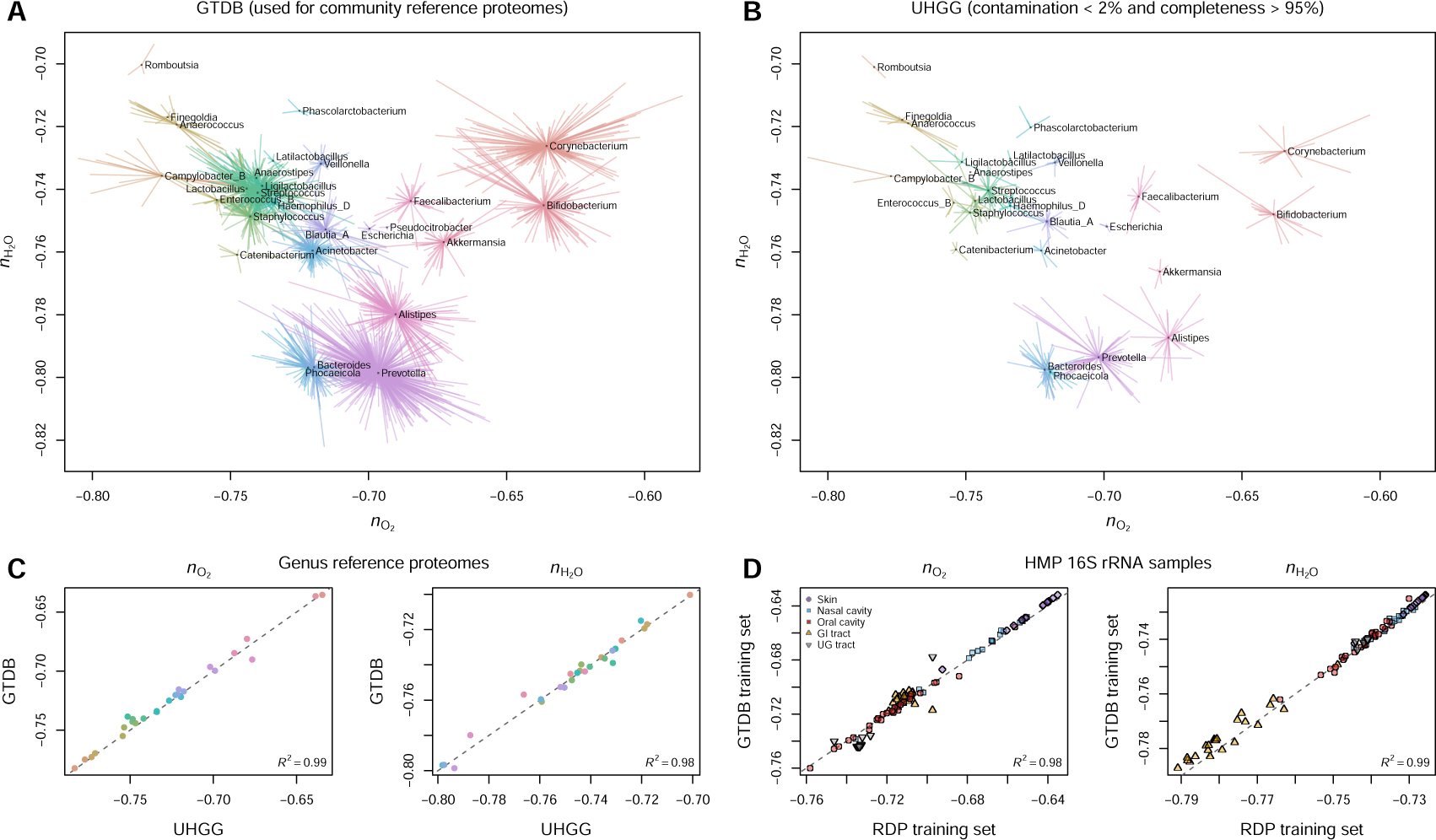
Comparison of reference databases for contamination assessment. (**A**) Chemical metrics of reference proteomes for selected genera and their species in GTDB; no contamination filtering was applied. The genera are those with a relative abundance difference of at least 5% between aggregated samples for controls and patients in one or more COVID-19 or IBD 16S rRNA datasets (see Section Differential contributions by obligate anaerobes and facultative anaerobes). The starburst patterns are drawn from a central point representing the parent taxon (genus) to each of the children (species). (**B**) Chemical metrics of reference proteomes for the same genera in the UHGG computed from species-levels clusters with contamination < 2% and completeness > 95%. (**C**) Comparison of chemical metrics for genus reference proteomes in GTDB and UHGG. (**D**) Comparison of chemical metrics of community reference proteomes for HMP samples generated with the GTDB-based training set used in this study and with the RDP training set used previously by Dick and Tan (2023). Dashed lines in (C) and (D) are 1:1 lines.

### Chemical variation of the human microbiome inferred from multi-omics data for body sites

The dataset of 16S rRNA gene sequences reported by Boix-Amorós et al (2021) represents nasal, skin, oral, and fecal (i.e., gut) communities. Reference proteomes for oral and nasal communities exhibit the lowest and highest ranges of *n*_O2_ compared to other body sites. Skin and gut communities have intermediate *n*_O2_ whereas gut communities have lower *n*_H2 O_ than other body sites (Figure 3A). Furthermore, Boix-Amorós et al (2021) performed viral inactivation experiments by treating samples with ethanol, formaldehyde, heat, psoralen, or trizol. Treatment with ethanol, and to a lesser extent heat or trizol, results in lower *n*_H2 O_ of community reference proteomes; no systematic change in *n*_O2_ was observed here (Figure 3–figure supplement 1).

**Fig. 3.**
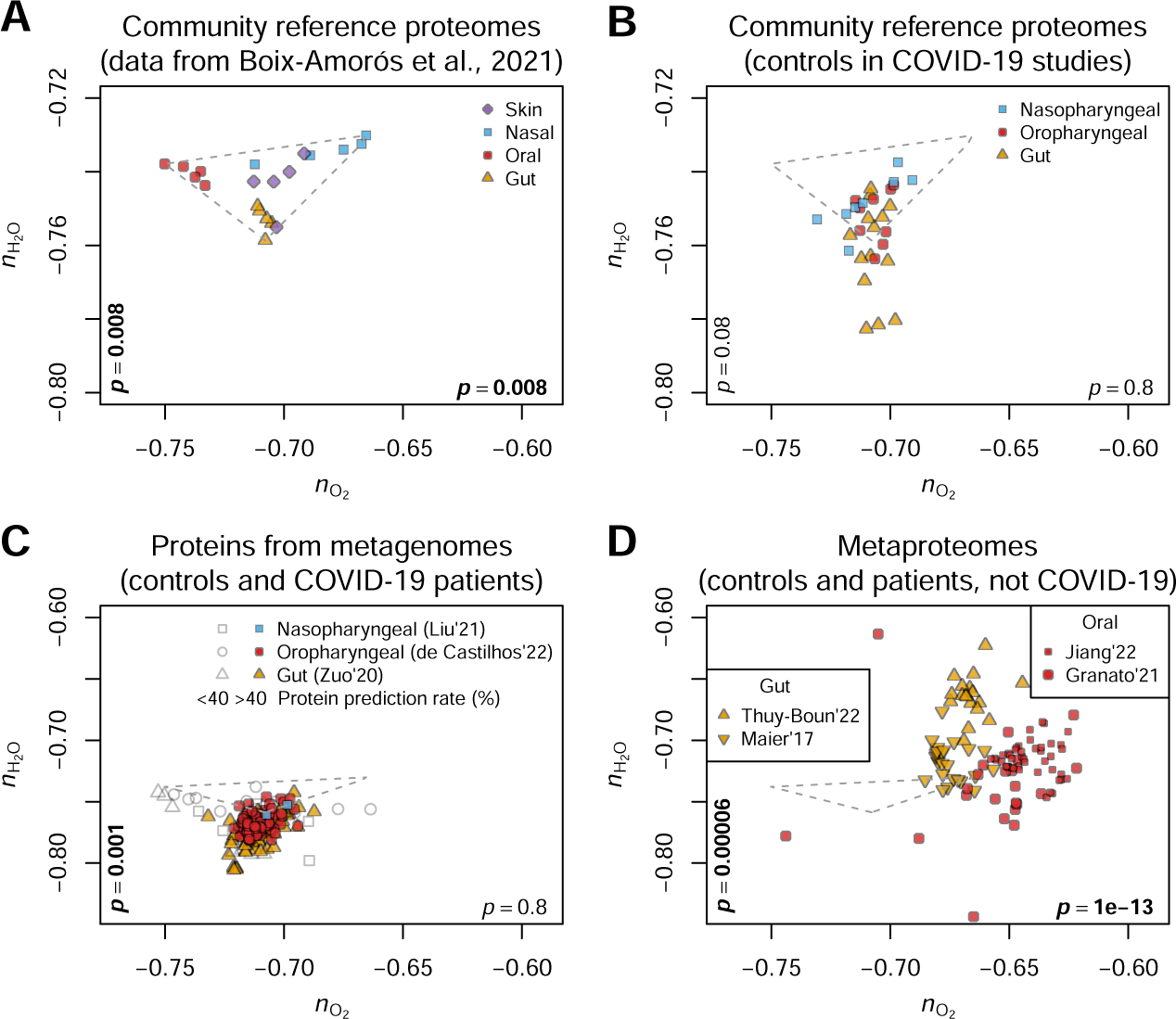
Multi-omics comparison of chemical metrics for microbiomes in human body sites. (**A**) *n*_O2_ and *n*_H2O_ of community reference proteomes for nasal, oral, skin, and gut sites, based on 16S rRNA gene sequences from Boix-Amorós et al (2021). (**B**) Community reference proteomes based on 16S rRNA data for controls in COVID-19 studies (Table 1). (**C**) Proteins predicted from metagenomes for controls and COVID-19 patients: nasopharyngeal (Liu et al, 2021), oropharyngeal (de Castilhos et al, 2022), and gut (Zuo et al, 2020). (**D**) Metaproteomes for controls and patients in non-COVID-19 studies: ulcerative colitis (Thuy-Boun et al, 2022) and dietary resistant starch (Maier et al, 2017) for gut microbiomes and oral cancer (Granato et al, 2021) and lung cancer (Jiang et al, 2022) for oral microbiomes. Each point represents a single sample, except for (B), where each point represents the mean of samples in a particular dataset. For visual comparison, the dashed triangle representing the convex hull around the points in (A) is replicated in (B–D). *p*-values (Wilcoxon unpaired test; bold text indicates *p* < 0.05) shown on the horizontal and vertical axes were calculated for *n*_O2_ and *n*_H2 O_, respectively, for gut samples compared to oral or oropharyngeal samples. **Figure supplement 1.** Differences of *n*_O2_ and *n*_H2 O_ between untreated and viral-inactivated samples.

**Table 1.** Sources of 16S rRNA gene sequence data for COVID-19 and IBD. Accessions are NCBI BioProject numbers. Numbers in the left column correspond to plotting symbols in Figs. 4 and 6. Sample counts exclude samples with low taxonomic classification rate (see Methods); datasets in each category are ordered by increasing geometric mean of numbers of control and patient samples. Asterisks (*) indicate datasets for mucosal samples; all other gut community data are for fecal samples. Several studies (Gevers et al, 2014; Lo Presti et al, 2019; Weng et al, 2019) reported data for both mucosal and fecal samples; only the data for fecal samples were analyzed here.

| No. | Accession and Reference | $n_{\text{Control}}$ | $n_{\text{Patient}}$ | No. | Accession and Reference | $n_{\text{Control}}$ | $n_{\text{Patient}}$ |
| --- | --- | --- | --- | --- | --- | --- | --- |
| Nasopharyngeal (COVID-19) |  |  |  | Oral or oropharyngeal (COVID-19) |  |  |  |
| 1 | PRJNA774098 Prasad et al (2022) | 12 | 21 | 1 | PRJNA767939 Rafiqul Islam et al (2022) | 15 | 22 |
| 2 | PRJNA714242 Smith et al (2021) | 10 | 32 | 2 | PRJNA692359 Iebba et al (2021) | 14 | 25 |
| 3 | PRJNA726205 Hernández-Terán et al (2021) | 7 | 76 | 3 | PRJNA780671 Gupta et al (2022a) | 24 | 30 |
| 4 | PRJNA673585 Ventero et al (2021) | 18 | 56 | 4 | PRJNA684070 Wu et al (2021) | 44 | 50 |
| 5 | PRJNA726992 and PRJNA726994 Shilts et al (2022) | 20 | 75 | 5 | PRJNA639286 Xu et al (2021) | 24 | 97 |
| 6 | PRJNA777915 Crovetto et al (2022) | 38 | 38 | 6 | PRJNA669421 Miller et al (2021) | 54 | 46 |
| 7 | PRJNA707350 Gupta et al (2022b) | 26 | 63 | 7 | PRJNA683617 Merenstein et al (2021) | 33 | 220 |
| 8 | PRJNA683617 Merenstein et al (2021) | 34 | 215 | 8 | PRJNA739539 Gao et al (2021) | 140 | 166 |
|  |  |  |  | 9 | PRJNA660302 Ren et al (2021) | 150 | 242 |
| Gut (COVID-19) |  |  |  | Gut (IBD) |  |  |  |
| 1 | PRJNA736160 Zhou et al (2021) | 14 | 15 | 1 | PRJNA679275 | 10 | 10 |
| 2 | PRJNA767939 Rafiqul Islam et al (2022) | 15 | 22 |  | Zakerska-Banaszak et al (2021) |  |  |
| 3 | PRJNA705797 Khan et al (2021) | 14 | 25 | 2 | PRJNA673073 Al-Amrah et al (2023) | 10 | 11 |
| 4 | PRJNA684070 Wu et al (2021) | 32 | 17 | 3 | PRJNA884507 Dahal et al (2023) | 10 | 11 |
| 5 | PRJNA703303 Chen et al (2022) | 30 | 26 | 4 | PRJEB33711 Alam et al (2020) | 9 | 21 |
| 6 | PRJNA636824 Gu et al (2020) | 30 | 30 | 5 | PRJNA975689 Park et al (2023) | 6 | 42 |
| 7 | PRJNA678695 Newsome et al (2021) | 34 | 50 | 6 | PRJNA313074 Mar et al (2016) | 13 | 30 |
| 8 | PRJNA753792 Romani et al (2022) | 22 | 89 | 7 | PRJNA391149 Lo Presti et al (2019) | 22 | 19 |
| 9 | PRJDB11949 and PRJDB12349 Mizutani et al (2022) | 61 | 45 | 8 | PRJNA368966 Bajer et al (2017) | 32 | 77 |
| 10 | PRJNA859805 Wang et al (2023) | 52 | 59 | 9 | PRJEB42155 Mills et al (2022) | 19 | 185 |
| 11 | PRJNA769052 Al Bataineh et al (2021) | 50 | 78 | 10 | PRJNA398089 | 43 | 112 |
| 12 | PRJNA758913 Ferreira-Junior et al (2022) | 38 | 141 |  | Lloyd-Price et al (2019) (*) |  |  |
| 13 | PRJNA660302 Ren et al (2021) | 72 | 100 | 11 | PRJEB47161 and PRJEB47162 Rausch et al (2023) | 96 | 56 |
| 14 | PRJNA756849 Schult et al (2022) | 145 | 104 | 12 | PRJEB18471 Halfvarson et al (2017) | 59 | 114 |
|  |  |  |  | 13 | PRJNA237362 Gevers et al (2014) | 31 | 224 |
|  |  |  |  | 14 | PRJNA431126 Weng et al (2019) | 38 | 286 |
|  |  |  |  | 15 | PRJNA398187 Ryan et al (2020) (*) | 63 | 282 |

To find out whether differences between body sites are recapitulated in independent datasets, chemical metrics were calculated for community reference proteomes for control subjects in COVID-19 studies (Table 1). Some of the datasets for gut communities exhibit relatively low *n*_H2O_, as indicated by their position below the triangular area that circumscribes values for the Boix-Amorós et al. dataset (Figure 3B). Turning to a multi-omics analysis, metagenomes for controls and COVID-19 patients have protein sequences that also exhibit overall lower *n*_H2O_ of gut compared to oral communities (Figure 3C). Finally, metaproteomes for gut communities have significantly lower *n*_O2_ and higher *n*_H2 O_ than those from oral communities (Figure 3D). In summary, community reference proteomes and metagenomes both reveal generally lower *n*_H2O_ for gut communities compared to other body sites, whereas metaproteomes of gut communities are reduced compared to oral communities. Rather than posing a conflict, the contrasting trends reflect different timescales of evolutionary and cellular processes: metagenomes and community reference proteomes probe genomic differences that arise through evolution, whereas metaproteomes reflect not only genomic constraints but also protein expression on cellular timescales.

### Chemical variation of the human microbiome inferred from multi-omics data for COVID-19 and IBD

Community reference proteomes for nasopharyngeal and oropharyngeal or oral samples in COVID-19 studies exhibit a range of positive and negative mean differences of chemical metrics between controls and patients (Figure 4A). Although the majority of datasets for oropharyngeal communities have lower mean *n*_O2_ in COVID-19 patients compared to controls, the overall difference for the datasets compiled here is not significant. In contrast, the overall difference for the compiled gut datasets is significant. Furthermore, the most extreme points correspond to large negative differences; six gut datasets have Δ*n*_O2_ < −0.005 but no gut dataset has Δ*n*_O2_ > 0.005.

**Fig. 4.**
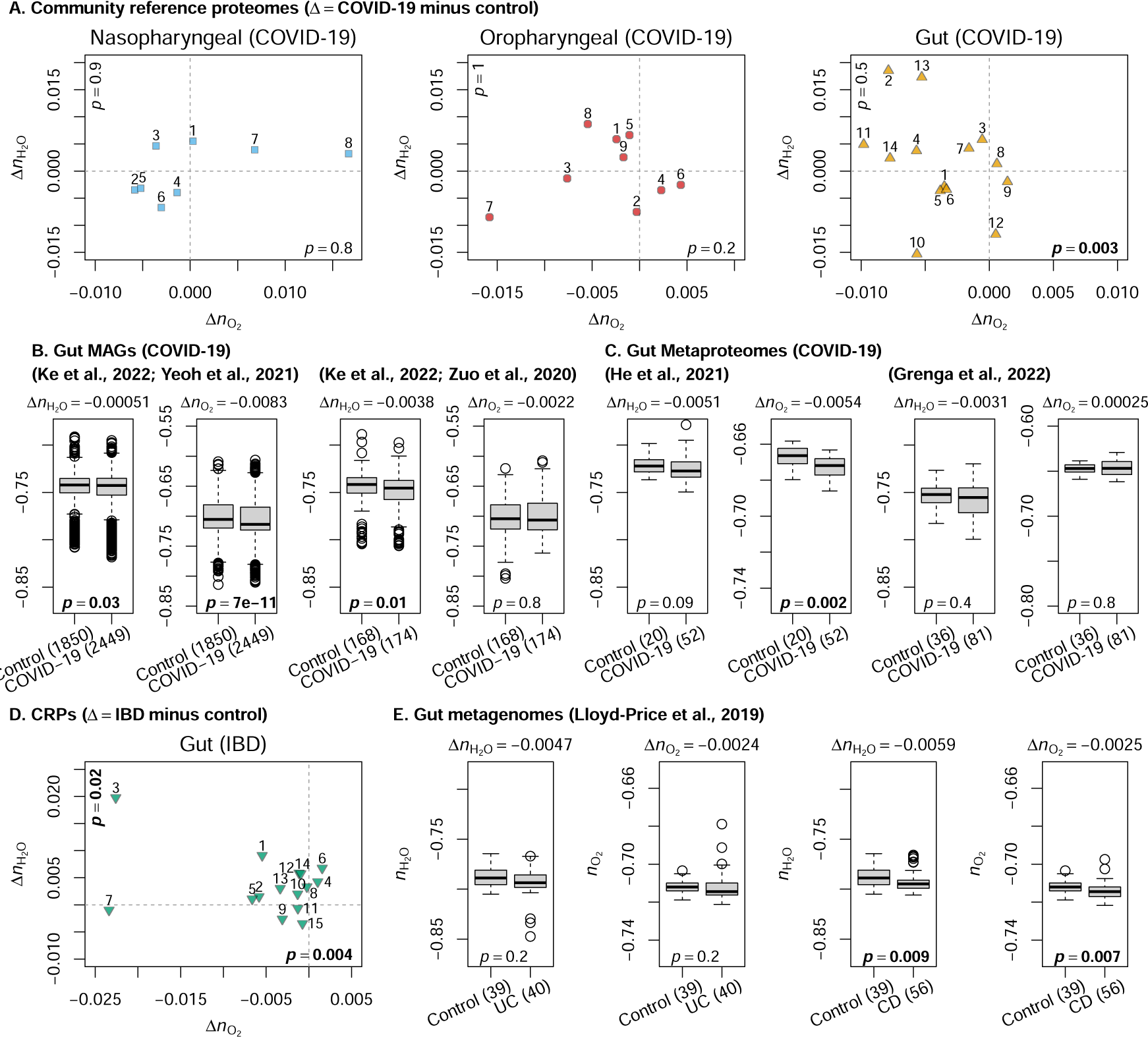
Differences of chemical metrics for microbial proteins between controls and COVID-19 or IBD patients. (**A**) Mean differences of chemical metrics for reference proteomes for nasopharyngeal, oropharyngeal, and gut communities from 16S rRNA datasets for COVID-19 studies (see Table 1). *p*-values (Wilcoxon paired test) are shown for differences of *n*_O2_ and *n*_H2O_ on the horizontal and vertical axes, respectively. (**B**) MAGs in controls and COVID-19 positive subjects reported by Ke et al (2022) based on metagenomic data from Yeoh et al (2021) and Zuo et al (2020). (**C**) Bacterial metaproteomes in controls and COVID-19 positive subjects based on data from He et al (2021) and Grenga et al (2022). (**D**) Community reference proteomes computed from the 16S rRNA datasets for IBD listed in Table 1. (**E**) Metagenomes for ulcerative colitis (UC) and Crohn’s disease (CD) patients and controls from Lloyd-Price et al (2019). In (B), (C), and (E), *p*-values were calculated using the unpaired Wilcoxon rank-sum test; boxplots show the median (thick line), first and third quartiles (box), most extreme values within 1.5 × the interquartile range away from the box (whiskers), and outliers (points); median differences of *n*_O2_ or *n*_H2 O_ are listed above the boxplots.

**Fig. 5.**
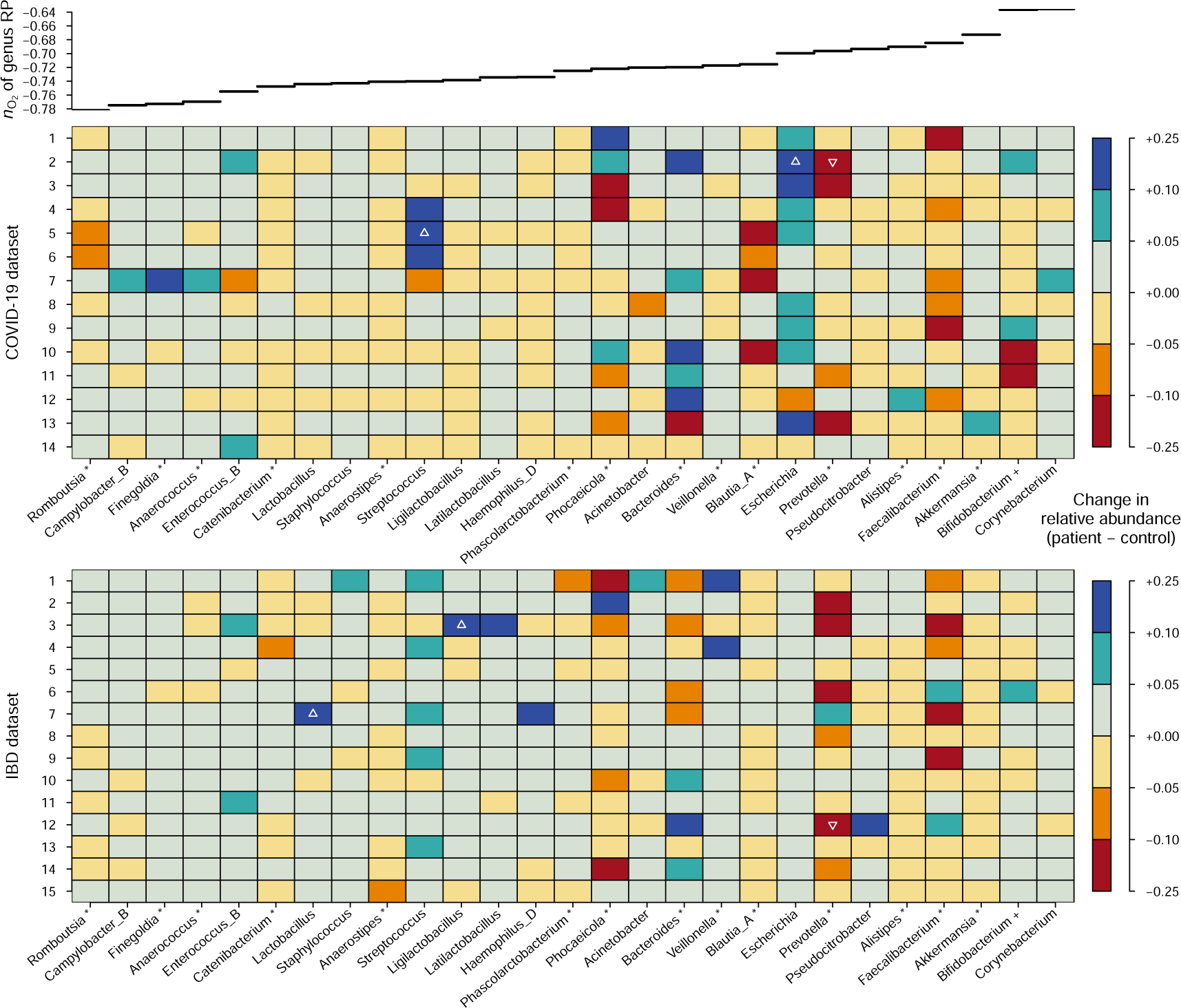
Differences of relative abundance of genera between aggregated samples for controls and patients in COVID-19 and IBD datasets. Genus-level classifications for samples in each dataset were summed to obtain aggregate counts for groups of control and patient samples, which were then divided by the total number of genus-level classifications in each group to obtain relative abundances. Genera with a relative abundance change of at least 0.05 (i.e., 5%) between controls and patients in any COVID-19 or IBD dataset are shown in the plot. Blue and red represent increased and decreased relative abundance, respectively, in patients compared to controls. The most intense colors represent a maximum relative abundance difference of 25%; differences larger than this are indicated by upor down-pointing triangles for increased or decreased abundances, respectively. The genera are ordered by increasing *n*_O2_ of their reference proteomes, as shown in the upper line plot. Obligately anaerobic genera in the list modified from Million and Raoult (2018) (see Methods) are indicated by an asterisk (*). Bifidobacterium, which comprises both obligately anaerobic and aerotolerant species and is classified as “variable” in this list, is indicated with a plus sign (+). **Figure supplement 1.** Chemical metrics of reference proteomes for genera with known oxygen tolerance.

This chemical reduction trend is supported by multi-omics data. Ke et al (2022) reported metagenome-assembled genomes (MAGs) that are in turn based on two previous metagenomic studies. The MAGs based on data from Yeoh et al (2021) and Zuo et al (2020) are characterized by lower median *n*_O2_ for protein sequences in COVID-19 compared to controls; however, only the difference for the former set of MAGs is significant (Figure 4B). Similarly, metaproteomic data reported by He et al (2021) yield significantly lower *n*_O2_ for bacterial proteins in COVID-19 patients than controls, but the metaproteomic dataset of Grenga et al (2022) shows no significant difference of chemical metrics for bacterial proteins (Figure 4C).

To get more information about the effects of inflammatory conditions on genomic adaptation, 16S rRNA and metagenomic data for IBD were also analyzed. In the compilation of 16S rRNA datasets, community reference proteomes have significantly lower *n*_O2_ in IBD compared to controls (Figure 4D). Analysis of shotgun metagenomic data from Lloyd-Price et al (2019) also yields lower *n*_O2_ of proteins in IBD compared to controls, but the difference is more significant for Crohn’s disease compared to ulcerative colitis (Figure 4E).

In summary, a large majority of 16S rRNA-based community reference proteomes point to lower stoichiometric oxidation state of gut microbial proteins in COVID-19 and IBD patients than in controls. Not all MAGs, metagenomes, and metaproteomes show significant differences of protein *n*_O2_ , but when they do, they recapitulate the chemical reduction trend in patients that was initially identified using community reference proteomes.

### Differential contributions by obligate anaerobes and facultative anaerobes

Dissecting the contributions of obligate anaerobes and aerotolerant organisms can help to understand the community-level trends of *n*_O2_ for COVID-19 and IBD. A list of genera with known oxygen tolerance was obtained from Million and Raoult (2018) and modified as described in the Methods. As a group, reference proteomes of aerotolerant organisms in this list have significantly higher *n*_O2_ compared to obligate anaerobes (Figure 5–figure supplement 1), indicating genomic adaptation to oxygen availability in a broad environmental context. “Obligate anaerobe” is a widely used term that hides a great deal of variation in actual oxygen tolerance. The functional definition of obligate anaerobes – organisms that do not require O_2_ to grow well and whose growth is blocked by exposure to a certain level of O_2_ in laboratory tests – means that many such organisms can survive or even grow at low oxygen levels (Lu and Imlay, 2021). A notable example is Bacteroides, some species of which can tolerate exposure to air for 24 h or more, yet this genus, or the Bacteroidia class, is commonly described as obligately anaerobic (Lu and Imlay, 2021; Winter and Bäumler, 2023).

Differences of relative abundance of genera between aggregated samples for controls and COVID-19 or IBD patients are visualized in Figure 5. Only genera with at least a 5% increase or decrease in at least one dataset are shown. Relative abundance differences rather than fold changes are used to visualize taxa with the largest abundance differences. For example, a genus with a 2-fold change from 10% to 20% relative abundance would be shown here, but one with a 2-fold change from 1% to 2% relative abundance would not be shown. *Faecalibacterium* and *Prevotella*, both obligate anaerobes, are notable for large relative abundance decreases in many datasets for COVID-19 and IBD. Interestingly, the relative abundance of *Blautia_A* (an obligately anaerobic genus) decreases by more than 5% and that of *Escherichia* (an aerotolerant genus) increases by more than 5% in many COVID-19 datasets, in contrast to differences of less than 5% in all the IBD datasets. The relative abundance of the aerotolerant *Streptococcus* increases by at least 5% in several datasets for both COVID-19 and IBD. *Bacteroides*, classified as an obligate anaerobe, is characterized by increased relative abundance in most COVID-19 datasets but has a more variable pattern in IBD; conversely, the obligate anaerobe *Phocaeicola* shows lower abundance in most IBD datasets but is more variable in COVID-19.

Despite being obligate anaerobes, important gut bacteria such as *Faecalibacterium* and *Prevotella* have relatively high *n*_O2_ of their reference proteomes, as shown by the upper line plot in Figure 5. Because of different taxonomic composition, the subcommunities of obligate anaerobes in fecal samples have higher abundance-weighted *n*_O2_ compared to subcommunities of obligate anaerobes in other body sites (Figure 6A). This trend underlies an interesting inversion, in which subcommunities of obligate anaerobes in the gut actually have more oxidized reference proteomes (i.e., have higher *n*_O2_ ) than subcommunities of aerotolerant organisms, as shown in Figure 6B for selected selected datasets for COVID-19 and IBD. Among all 16S rRNA datasets in the compilation, this inversion characterizes the gut more so than other body sites (Figure 6–figure supplement 1).

**Fig. 6.**
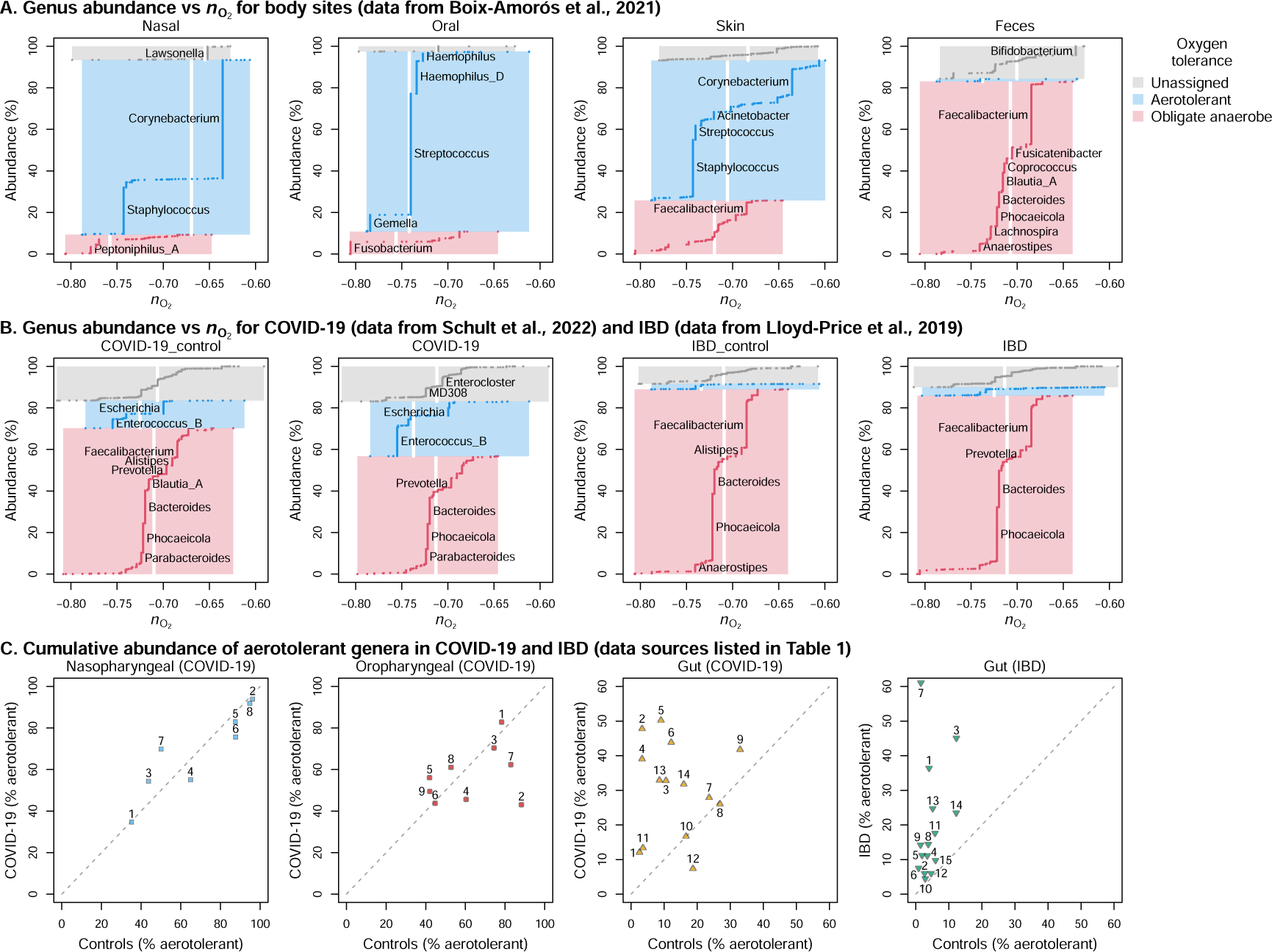
Obligate anaerobes and aerotolerant genera vary in abundance and have different stoichiometric oxidation states of proteins depending on body sites. In (**A**) for body sites and (**B**) for gut communities in selected COVID-19 and IBD studies, vertical lines denote relative abundances of genera across all samples for each body site or clinical condition and colors represent aerotolerance groups. Genera that are more than 3% abundant are labeled. Within aerotolerance groups, lines are arranged by increasing *n*_O2_ of the reference proteomes for genera, and the width and height of shaded areas represent the total range of *n*_O2_ and cumulative abundance. Vertical white lines indicate abundance-weighted mean values of *n*_O2_ for genera in each group. (**C**) Cumulative abundance of aerotolerant genera as a percentage of all aerotolerant and obligately anaerobic genera in COVID-19 and IBD studies. Points that plot above the diagonal dashed 1:1 line represent datasets with higher abundance of aerotolerant genera in patient compared to control groups. See Table 1 for key to datasets. **Figure supplement 1.** Differences of *n*_O2_ and *n*_H2O_ between subcommunities of obligate anaerobes and aerotolerant genera in controls and patients.

Gut microbiota but not nasopharyngeal or oropharyngeal microbiota are frequently enriched in aerotolerant genera in COVID-19 or IBD patients compared to controls (Figure 6C). The highest percentage of aerotolerant genera for patient groups in any gut dataset (up to ca. 50% and 60% for COVID-19 and IBD, respectively) is lower than the maximum observed for nasopharyngeal or oropharyngeal samples, reflecting the persistence of obligate anaerobes in the gut even in disease conditions. The widely varying percentages of aerotolerant genera among datasets for either COVID-19 and IBD could indicate increased oxygen levels in the gut for most, but not all, patients represented by this data compilation. While an expansion of aerotolerant gut microbes is consistent with previous reports for inflammatory diseases (Shin et al, 2015; Dubourg et al, 2016), the chemical reduction trend (i.e. lower *n*_O2_ of proteins) in multi-omics datasets for gut communities in COVID-19 and IBD patients compared to controls is a surprising result that evokes not only environmental factors but also competitive interactions.

## Discussion

Chemical variables play important roles for host-associated as well as for free-living microbiomes (Lee et al, 2022; Pfister et al, 2022). Oxidation/reduction and hydration/dehydration processes contribute fundamentally to metabolism and host physiology, so this study focused on the calculation of corresponding chemical metrics for protein sequences inferred from multi-omics datasets to document community-level chemical differences (summarized in Table 2). The unexpected findings include a signature of dehydration for gut communities compared to other body sites and a shift to more reduced proteins of gut communities in COVID-19 and IBD patients compared to controls.

**Table 2.** Most significant differences of chemical metrics in this study. CRP – community reference proteome; MAG – metagenome-assembled genome; MG – metagenome; MP – metaproteome.

| Comparison | Difference | Data Type | Figure |
| --- | --- | --- | --- |
| Nasal vs other sites | $\uparrow n_{O_2}$ | CRP | 3A |
| Oral vs other sites | $\downarrow n_{O_2}$ | CRP | 3A |
| Gut vs other sites | $\downarrow n_{H_2O}$ | CRP | 3A |
| Gut vs oral | $\downarrow n_{H_2O}$ | MG | 3C |
| Gut vs oral | $\downarrow n_{O_2}$ | MP | 3D |
| Ethanol treatment vs untreated | $\downarrow n_{H_2O}$ | CRP | 3–supp. 1 |
| Gut COVID-19 vs control | $\downarrow n_{O_2}$ | CRP | 4A |
| Gut COVID-19 vs control | $\downarrow n_{O_2}$ | MAG | 4B |
| Gut COVID-19 vs control | $\downarrow n_{O_2}$ | MP | 4C |
| Gut IBD vs control | $\downarrow n_{O_2}$ | CRP | 4D |
| Gut IBD vs control | $\downarrow n_{O_2}$ | MG | 4F |

Community reference proteomes for skin and nasal communities are generally more oxidized than those for oral communities. This pattern parallels previous analyses of metagenomic data from the HMP (Vecchio-Pagan et al, 2017; Dick and Tan, 2023) and suggests that oxygenated habitats on the skin and in the airway select for genomes that code for relatively oxidized proteins. Community reference proteomes are more dehydrated for gut communities than for other body sites and are more dehydrated for samples treated with ethanol than for untreated samples. The intestine has a physiological function of water absorption (Leiper, 2015), and the dehydrating effect of ethanol is described in protocols for preparing tissue sections for histological analysis (Lai and Lü, 2012). The results therefore support the notion of community-level genomic adaptation to dehydrating forces that microbes may encounter in the gut or other environments.

Turning to a multi-omics analysis, community reference proteomes and proteins predicted from metagenomes show similar ranges of *n*_O2_ and *n*_H2 O_, but metaproteomes are relatively oxidized. A higher abundance of cytoplasmic than membrane proteins, as well as greater extraction efficiency for cytoplasmic proteins in metaproteomic experiments, can explain the relatively high oxidation state for metaproteomes (Dick and Meng, 2023). Interestingly, gut metaproteomes are more reduced than oral ones, which is the opposite trend from community reference proteomes. It therefore appears that low oxygen concentrations in the intestinal lumen (e.g., Ast and Mootha, 2019; Pfister et al, 2022) may have a more pronounced effect on protein expression, as detected by metaproteomes, than on genomic adaptation.

Multi-omics datasets for COVID-19 and community reference proteomes for IBD are characterized by relatively reduced proteins in disease. Because oxidizing conditions are thermodynamically predicted to select for relatively oxidized rather than reduced proteins (Dick and Meng, 2023), this signature of chemical reduction at first appears incompatible with the higher oxygen levels and/or greater availability of other electron acceptors previously associated with inflammation (Rigottier-Gois, 2013; Pfister et al, 2022; Winter and Bäumler, 2023). The value of a thermodynamic model, as a kind of null theory, is that the failure of a specific thermodynamic prediction indicates the existence of mechanisms that are more complex than the simple model assumes (Harte, 2004). If oxidizing conditions are indeed characteristic of inflammation, the inference of relatively reduced proteins needs to be explained by some other means than thermodynamic control at the whole-community level. Natural O_2_ gradients within the intestine support populations of obligate anaerobes and facultative anaerobes in distinct locations (Donaldson et al, 2016). When inflammation occurs and conditions become more oxidizing, facultative anaerobes can be expected to increase in abundance. Indeed, this study found increased abundances of aerotolerant genera for patient compared to control groups in most COVID-19 and IBD datasets, which recapitulates previous findings for inflammatory diseases (e.g., Shin et al, 2015). This is accompanied by a lower abundance of obligate anaerobes; crucially, however, subcommunities of anaerobes in the gut actually have higher protein *n*_O2_ than subcommunities of aerotolerant organisms. This accounts for the chemical reduction trend at the whole-community level in patients and provides a record of genomic adaptation of obligate anaerobes to transiently higher oxygen levels associated with gut inflammation.

This study has several notable limitations. Although the gut is characterized by spatial gradients of oxygen concentration and water content, the analysis here of data generated from mostly fecal samples does not permit resolving community composition along spatial gradients. Mucous becomes denser and more continuous toward the rectum and may be associated with a longitudinal gradient of decreasing water content within the gut (Tropini et al, 2017); location-specific data would be useful to further test the hypothesized link between *n*_H2O_ of proteins and environmental hydration conditions. Although comparisons made above suggest that the numerical results are not greatly affected by contamination of available metagenomic data or genomic reference databases, additional checks such as analysis of laboratory blanks or in silico contaminant removal were not performed here. The similar differences of chemical metrics between sample groups for many 16S rRNA datasets as well as metagenomes or MAGs imply that the results described here are not artifacts of particular sampling or sequencing protocols.

## Conclusion

Through a stoichiometric analysis in which O_2_ and H_2_O are defined as thermodynamic components, it was shown that metagenomes and community reference proteomes in the gut code for proteins with lower *n*_H2O_ than other body sites. This is most easily explained as selection for genomes with relatively low hydration state in the context of physiological function of water absorption in the intestines. In contrast, metaproteomic data indicate that microbial protein expression in the gut is shifted toward more reduced proteins (those with low *n*_O2_ ) compared to oral communities. This leads to the hypothesis that the dynamics of genomic adaptation and protein expression, which occur on relatively long and short timescales, show differential sensitivity to hydration/dehydration and oxidation/reduction conditions. Protein expression experiments conducted under controlled oxidation and hydration conditions and analysis of chemical variation of genomes within multiple phylogenetic lineages with known habitat preference are some other approaches that could be used to test this hypothesis.

It was previously noted that metagenomes of nasal and skin microbiota have proteins that are relatively oxidized, presumably as a consequence of exposure to higher oxygen levels compared to oral and gut habitats (Vecchio-Pagan et al, 2017). This study identifies similar patterns in community reference proteomes but adds the initially puzzling observation that oral, rather than gut communities, have the most reduced protein sequences. Why gut communities are not relatively reduced, despite inhabiting a nominally reducing environment, is a question that leads to new insight. In particular, *Faecalibacterium*, an obligate anaerobe recognized as having anti-inflammatory associations (Sokol et al, 2008), has a reference proteome that is more oxidized than many anaerobes in other body sites and even than many aerotolerant bacteria in the gut. The lower abundance of *Faecalibacterium* in many datasets for COVID-19 and IBD patients compared to controls contributes to a trend of chemical reduction (i.e. lower *n*_O2_ ) associated with community reference proteomes for these diseases. This reduction trend is further supported by metagenomic and metaproteomic data. Synthesizing the available evidence, the relatively high oxidation state of obligate anaerobes in the gut represents a genomic adaptation to the oxidative environment of gut inflammation. An important implication is that anaerobes with highly reduced proteomes would be less well adapted to oxidative conditions in the intestine and therefore might be predicted to negatively influence a patient’s response to inflammatory disease.

## Methods

### Theory and calculation of chemical metrics

Concepts borrowed from geochemical thermodynamics provide a theoretical framework for linking elemental stoichiometry of biomolecules and the environment. A key concept is thermodynamic components; this is a set of chemical species that is the minimum number needed to represent the range of elemental composition of molecules in a system under study. The system here refers to all possible protein sequences; because the 20 common proteinogenic amino acids are made up of five elements, that is the number of thermodynamic components that is needed. Thermodynamic theory places no restriction on the components as long as they are mathematically feasible, so other considerations determine the utility of any possible set of components. As noted above, water and oxygen are major variables in metabolic reactions, so they are among the components chosen here. The remaining components, namely glutamine, glutamic acid, and cysteine (QEC) were chosen for several reasons. 1) They represent nitrogen, carbon, and sulfur in biologically available forms. Amino acids rather than other simple biomolecules are a reasonable choice for representing the chemical variation among proteins. 2) Glutamine and glutamic acid have a relatively high degree of metabolic network connectivity; that is, they are central metabolites (Dick et al, 2020). 3) Compared to most other combinations of amino acids, QEC yields a strong positive correlation between stoichiometric oxidation state (the number of O_2_ calculated for a given protein, normalized by protein length) and average oxidation state of carbon (*Z*_C_) (Dick et al, 2020). This is an important consideration because carbon oxidation state is an electronegativity-based measure that is independent of the choice of components, and the close alignment between alternative metrics for oxidation state (*Z*_C_ and *n*_O2_ ) reflects an underlying consistency. 4) For all the proteins in a given genome, there is very little correlation between *Z*_C_ and *n*_H2O_ (i.e. stoichiometric hydration state) (Dick, 2022). This is important because it shows that metrics for oxidation and hydration state measure different aspects of the chemical variation between proteins.

In a chemical sense, thermodynamic components are a minimal set of species, or basis species, that can be used to balance a theoretical reaction between any two proteins. A limitation of the theory is that such a reaction does not represent the actual mechanism of protein formation; the inclusion of highly-connected metabolites as components mitigates this concern somewhat. Mathematically, thermodynamic components represent a transformation from elemental composition to chemical composition (Eq. 1). Thermodynamic components can also be conceptualized as a projection from the higher-dimensional compositional space of 20 amino acids. Unlike statistical dimensionality reduction techniques, thermodynamic components have a stable definition that doesn’t depend on the data being analyzed. This means that amino acid compositions of proteins inferred from multi-omics datasets can be used to test the hypothesis that oxygen and water availability shape the genomic adaptation of the human microbiota.

*n*_H2 O_ and *n*_O2_ were calculated according to

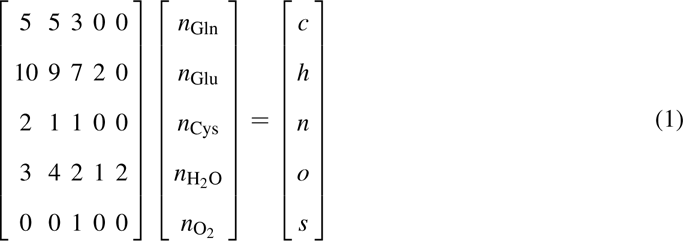

where *n*_Gln_, *n*_Glu_, *n*_Cys_, *n*_H2 O_, and *n*_O2_ are the stoichiometric coefficients of a theoretical reaction to form a protein with formula C*_c_*H*_h_*N*_n_*O*_o_*S*_s_*, and the stoichiometric matrix on the left represents the number of elements in each of the basis species; the values of *n*_H2 O_ and *n*_O2_ reported here were normalized by protein length. To facilitate the processing of multi-omics datasets, instead of calculating elemental formulas of proteins as an intermediate step, amino acid compositions were combined with precomputed values of *n*_O2_ and *n*_H2 O_ for amino acid residues to calculate the chemical metrics for proteins. The specific functions used for this are calc_metrics() in the chem16S R package and nH2O() and nO2() in the canprot R package (Dick and Kang, 2023; Dick, 2021).

### 16S rRNA gene sequence processing

Literature searches were used to locate publicly available 16S rRNA gene sequencing datasets. Gut microbiome datasets with at least 20 samples (total for controls and patients) available by the end of 2023 were included, while those for oropharyngeal and nasopharyngeal microbiomes available by the end of 2022 were included. Sequence data were downloaded from the National Center for Biotechnology Information (NCBI) Sequence Read Archive (SRA). FASTQ sequence files for paired-end sequences were merged using the “fastq_mergepairs” command of VSEARCH version 2.15.0 (Rognes et al, 2016). Quality filtering was done with maximum expected error rate of 0.005 and a sequence truncation length specific for each dataset (see Supplementary file 2 for details and sequence processing statistics). Singletons were removed and remaining sequences were subsampled at a depth of 10000. Reference-based chimera detection with the VSEARCH command “uchime_ref” was performed using the SILVA SSURef NR99 database version 138.1 (Quast et al, 2012). Taxonomic classification was performed using the RDP Classifier version 2.13 (Cole et al, 2014) retrained with the Genome Taxonomy Database (bac120_ssu_reps and ar53_ssu_reps in GTDB release 207) (Parks et al, 2022). Samples with less than 100 classified reads were excluded from the following analysis.

### Reference proteomes for taxa

Reference proteomes of archaeal and bacterial taxa were made as described previously (Dick and Tan, 2023), except that GTDB was used instead of the NCBI Reference Sequence database (RefSeq). Briefly, for each genome in GTDB, the amino acid composition of all proteins was summed and divided by the number of proteins. Then, amino acid compositions for all genomes in each genus were summed and divided by the number of genomes to generate the amino acid compositions of genus reference proteomes. Similarly, amino acid compositions for all genomes in each family, order, class, and phylum were summed and divided by the number of genomes to generate the amino acid compositions of reference proteomes for taxa at those ranks.

Reference proteomes for species and genera in the Unified Human Gastrointestinal Genome (UHGG) were made in an analogous fashion. Taxonomic lineages and contamination and completeness values for each of the 4744 species-level clusters present in UHGG version 2.0.1 were obtained from the MGnify website (https://www.ebi.ac.uk/metagenomics/browse/genomes, accessed on 2023-12-30); those with contamination < 2% and completeness > 95% were used to generate reference proteomes for comparison with GTDB.

### Community reference proteomes

The chem16S package version 1.0.0 (Dick and Kang, 2023) was used to analyze chemical metrics of community reference proteomes generated by combining taxonomic classifications made using the GTDB (see above) with GTDB-based reference proteomes for taxa. The lowest-level taxonomic classification, from genus to phylum, for each processed 16S rRNA gene sequence was retained. For each sample, the abundances of taxa were multiplied by the amino acid compositions of reference proteomes for taxa and summed to obtain the amino acid composition of the community reference proteome.

### Metagenomic data processing

#### Shotgun metagenomes

For each sequencing run, only forward sequences were used, adapter sequences were removed, sequences were dereplicated, human sequences were removed using bowtie2 version 2.5.0 (Langmead and Salzberg, 2012) with the GRCh38 reference database (https://genome-idx.s3.amazonaws.com/bt/GRCh38_noalt_ as.zip, accessed on 2022-11-17), rRNA sequences were removed using SortMeRNA version 2.1b (Kopylova et al, 2012), and partial protein sequences were predicted using FragGeneScan version 1.18 (Rho et al, 2010). The amino acid compositions of all protein sequences in each run were summed and used to calculate chemical metrics. Dataset accession numbers and processing statistics are listed in Supplementary file 1.

#### Metagenome-assembled genomes from COVID-19 patients and controls

Nucleotide sequences of metagenome-assembled genomes (MAGs) generated by Ke et al (2022), which are based on metagenomic data originally reported by Yeoh et al (2021) and Zuo et al (2020) (NCBI BioProjects PRJNA650244 and PRJNA624223, respectively), were obtained from the file MAG.zip (https://figshare.com/s/a426a12b463758ed6a54, accessed on 2022-10-27). The MAGs analyzed here meet completeness and contamination thresholds of *≥*50% and *≤*5%, respectively (Ke et al, 2022). Identifications of MAGs from COVID-19 patients and controls were obtained from BioSample metadata for NCBI BioProject PRJNA650244. For each MAG, protein sequences were predicted using Prodigal (Hyatt et al, 2010), and the amino acid compositions of all predicted proteins were summed and used to calculate chemical metrics.

### Metaproteomic data processing

Protein sequences were obtained from GenBank (accessed on 2022-09-06), UniProt (accessed on 202208-31), or the Human Oral Microbiome Database (HOMD) (Chen et al, 2010) version 9.15 (updated date: 2022-02-07; accessed on 2023-02-03). For processing UniProt IDs, the UniProt ID mapping tool (Huang et al, 2011) was used with the taxonomy filter set to include only bacterial sequences (in order to exclude human proteins), and amino acid composition was computed from canonical protein sequences. For processing HOMD IDs, sequence files in the PROKKA directory were used (Genomes Annotated with PROKKA 1.14.6).

For the following datasets, protein sequences were obtained from the UniProt database, and amino acid compositions of proteins identified in each sample were summed and weighted by abundances (where available) to obtain the amino acid composition of the metaproteome. (Maier et al, 2017) (gut starch diet): Metaproteomic abundances and UniProt IDs were obtained from https://zenodo.org/record/838741. Additional mapping to the UniParc database was used to retrieve obsolete sequences. (Thuy-Boun et al, 2022) (ulcerative colitis): Protein IDs were extracted from PeptideEvidence fields of *.mzid.gz files from accession PXD022433 listed on ProteomeXchange (Deutsch et al, 2020). IDs with “.” or “_” were omitted to retain UniProt IDs.

For the saliva microbiome from Granato et al (2021), Majority Protein IDs and LFQ intensity for saliva cells were taken from Table S9 of the source publication; data for saliva supernatant were not analyzed here. For each protein, the first Majority Protein ID was used to look up the protein sequence in HOMD. Organism ID SEQF2791 (*Selenomonas* sp. HMT 136) was not found in HOMD at the time of this study, so the second Majority Protein ID was used in this case. For the oral microbiome from Jiang et al (2022), Majority Protein IDs and LFQ intensity were taken from the proteinGroups.txt file available downloaded from PXD026727. The first Majority Protein ID was used for each protein, except for some organisms not found in HOMD at the time of this study (SEQF1058, SEQF3075, SEQF1068, SEQF1063, SEQF2480, SEQF2762, SEQF3069, and SEQF2791), for which the second Majority Protein ID was used. For both of these datasets, the amino acid composition of each identified protein was multiplied by its abundance measured by LFQ intensity and summed to obtain the amino acid composition of the metaproteome.

For the COVID-19 gut metaproteome from He et al (2021), the file “Table S2. Global metaproteome.xlsx” was downloaded from Supplementary Files on ResearchSquare (https://doi.org/10.21203/ rs.3.rs-208797/v1). UniProt IDs were extracted from the “Accession” column by keeping values starting with “tr|” or “sp|”. Amino acid compositions of the proteins were multiplied by spectral counts and summed to obtain the amino acid composition of the metaproteome. For the COVID-19 gut metaproteome from Grenga et al (2022), GenBank protein IDs were extracted from *.mzid.gz files from accession PXD024990 listed on ProteomeXchange, and the esearch command (part of the NCBI E-utilities; https://www.ncbi.nlm.nih.gov/books/NBK25501/) with the “-organism bacteria” option was used to download bacterial protein sequences.

### Oxygen tolerance of genera

The “List of Prokaryotes according to their Aerotolerant or Obligate Anaerobic Metabolism” was taken from Table S1 of Million and Raoult (2018). The list was modified in this study by the removal of *Photorhabdus*, which was listed in both categories, and the addition of obligately anaerobic genera *Anaerobutyricum*, *Phocaeicola*, and *Romboutsia*, according to their descriptions in Shetty et al (2018); Wang et al (2021); Ricaboni et al (2016). With these changes, 234 obligately anaerobic and 399 aerotolerant genera are listed. Alphabetic suffixes for polyphyletic groups in GTDB (Parks et al, 2018) were removed before matching genus names to this list. Any genus listed as “variable” or “unknown” or not present in the list was considered to have unassigned oxygen tolerance.

### Statistics

R (R Core Team, 2023) was used for data visualization and statistical analysis. Wilcoxon tests were used to calculate *p*-values. Tests were made for unpaired observations except where noted. *p* < 0.05 was considered significant.

## Additional information

### Funding

No funding was received for this work.

### Author contributions

JMD performed the research and wrote the manuscript.

### Author ORCID

Jeffrey M. Dick https://orcid.org/0000-0002-0687-5890

### Competing interests

The author declares that no competing interests exist.

### Data availability

The original contributions of this study are are available in the following repositories. Training files for the RDP Classifier generated from GTDB release 207 are archived at https://doi.org/10.5281/zenodo. 7633100. Analysis scripts, processed data files generated in this study, and functions to make the plots are in the “microhum” section of the JMDplots R package version 1.2.19, available at https://github. com/jedick/JMDplots and archived at https://doi.org/10.5281/zenodo.10686108. The compiled HTML version of the vignette that run the functions to make the plots is available at https://chnosz.net/JMDplots/ vignettes/microhum.html.

## Supplementary files

**Supplementary file 1.** Metagenome sequence processing statistics.

**Supplementary file 2.** 16S rRNA gene sequence processing statistics.

---

**Figure 1–figure supplement 1.**
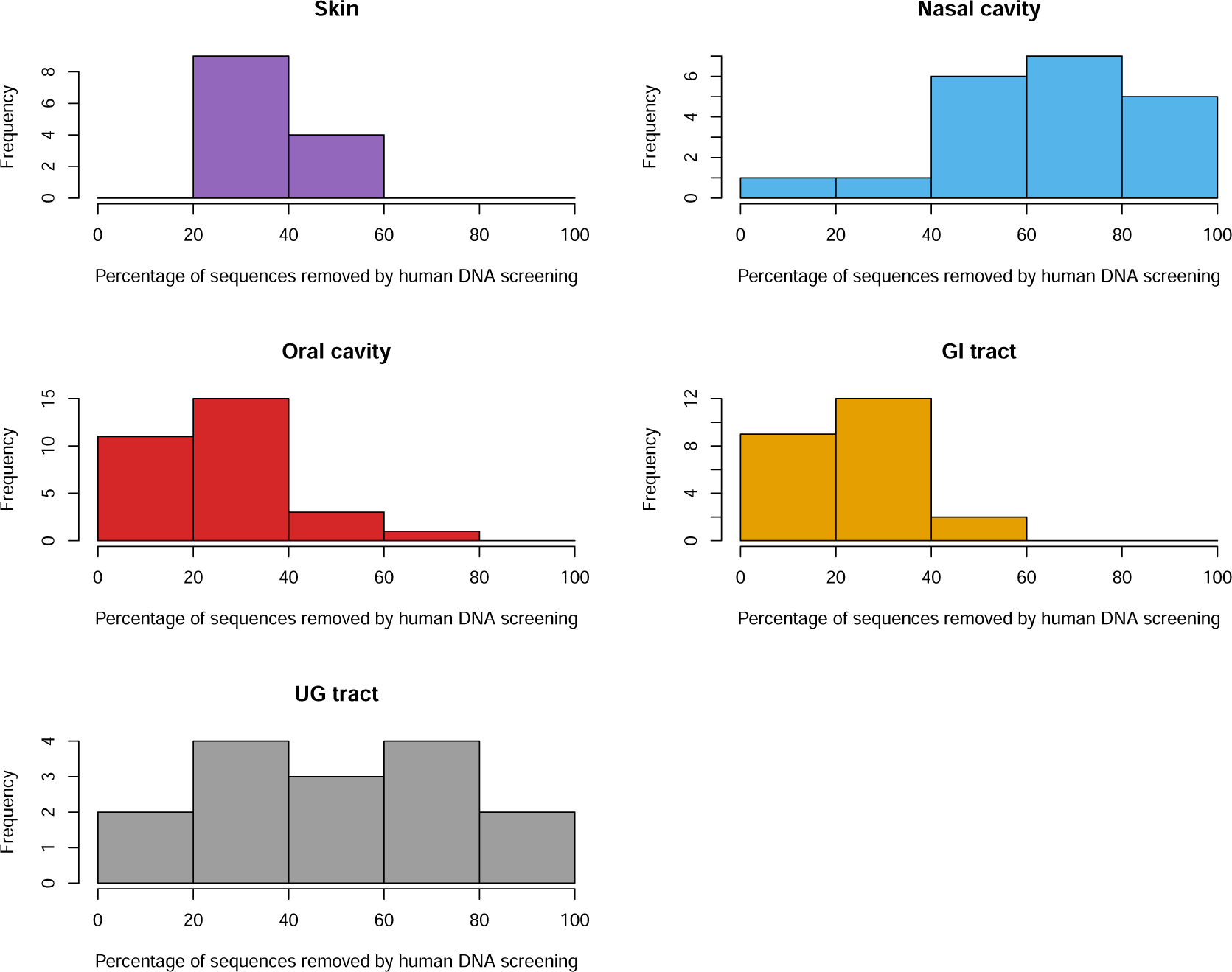
Percentages of sequences removed by human DNA screening of Human Microbiome Project metagenomes for various body sites.

**Figure 3–figure supplement 1.**
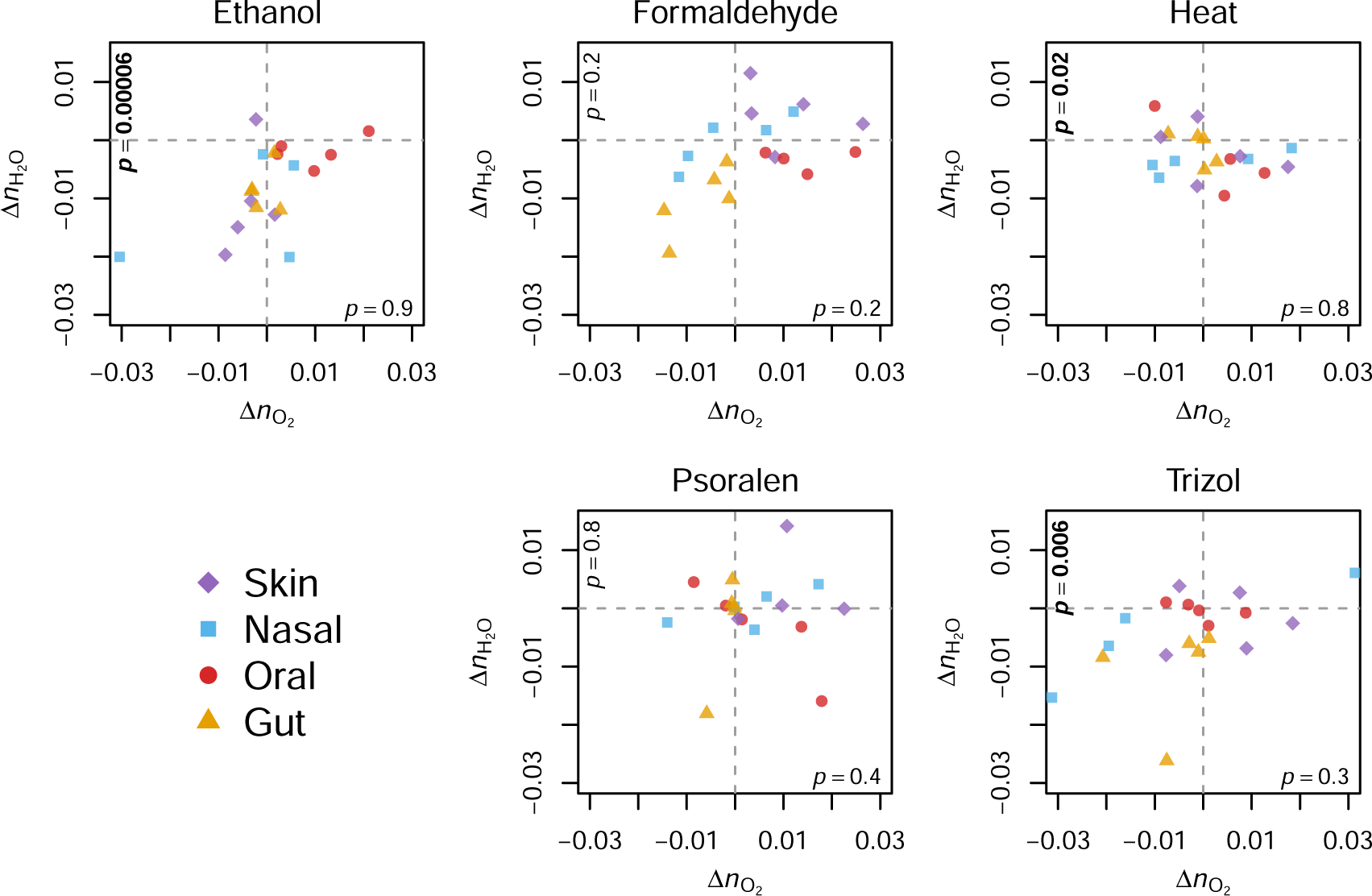
Differences of *n*_O2_ and *n*_H2 O_ between untreated and viral-inactivated samples. 16S rRNA gene sequence data were taken from Boix-Amorós et al (2021). Bold text indicates *p* < 0.05 (Wilcoxon paired test between untreated and viral-inactivated samples from the same subject).

**Figure 5–figure supplement 1.**
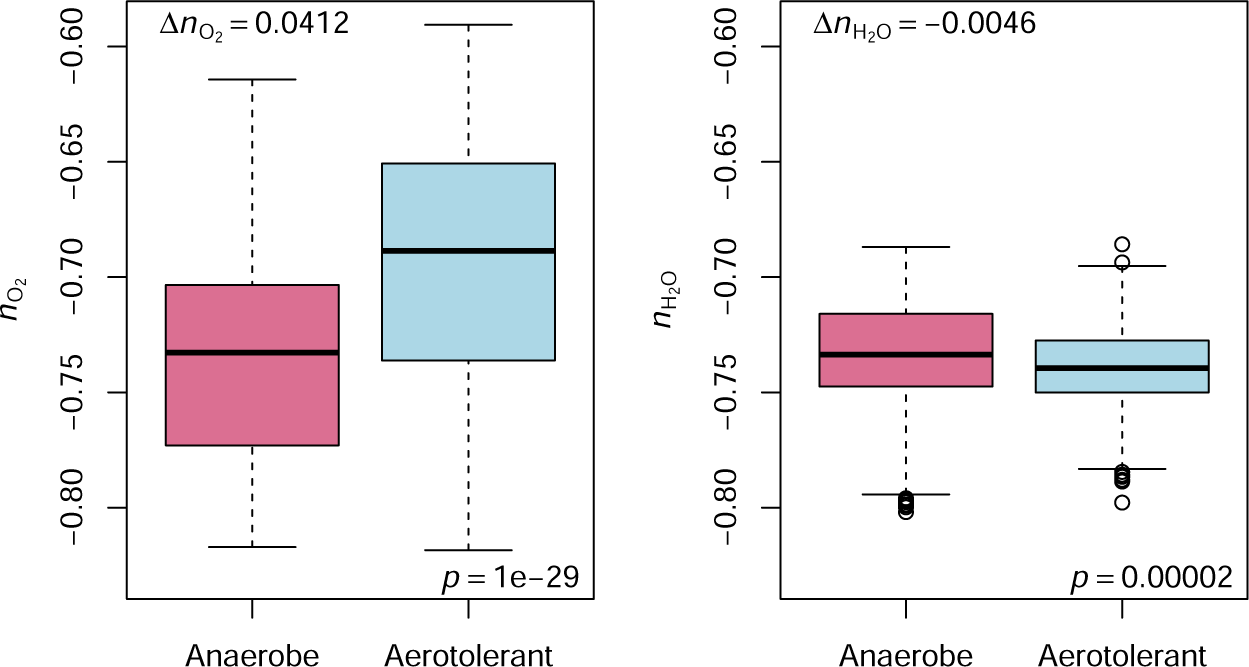
Chemical metrics of reference proteomes for genera with known oxygen tolerance. Boxplots show distribution of stoichiometric oxidation and hydration state (*n*_O2_ and *n*_H2 O_) for reference proteomes of prokaryotic genera assigned as obligately anaerobic or aerotolerant. Assignments were taken from Table S1 of Million and Raoult (2018) and modified as described in the Methods. Delta values denote mean differences of *n*_O2_ or *n*_H2 O_ between obligately anaerobic and aerotolerant genera.

**Figure 6–figure supplement 1.**
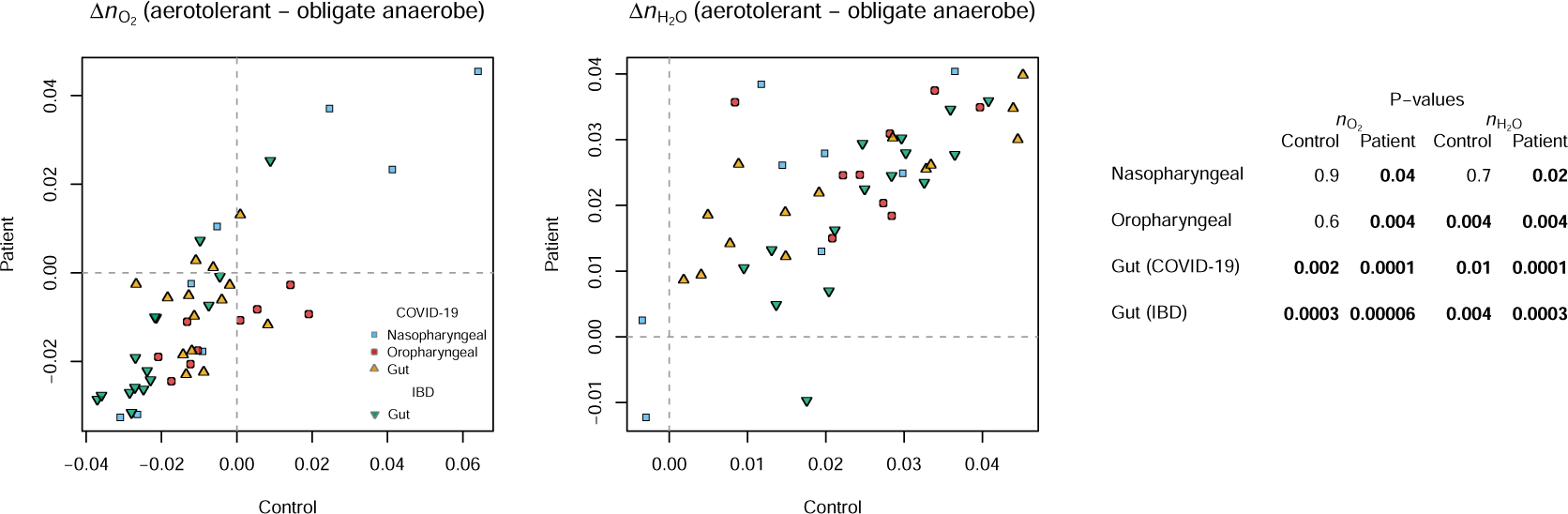
Differences of *n*_O2_ and *n*_H2 O_ between subcommunities of obligate anaerobes and aerotolerant genera in controls and patients.

## Notes

https://doi.org/10.5281/zenodo.10686108

### Competing Interest Statement

The authors have declared no competing interest.

### Summary of Updates

Major revision: Analyze HMP, UHGG, and IBD data and add Figures 1, 2, 5, and 6.

